# Effect of Emoji on Autonomic Nervous System: Evidence from EDA and RSA studies

**DOI:** 10.1101/2024.04.13.589339

**Authors:** Deeksha Patel, Abhinav Dixit, Om Lata Bhagat

## Abstract

**Background:** Facial expression, gesture, and posture play an important role in perceiving emotions during communication. In virtual communication platforms, users have devised and learned to use a variety of expressive emojis along with text messages to express certain emotions. The effect of emojis on human psychology and associated autonomic responses is still vague.

**Methods:** A total of 100 healthy individuals (50 males and 50 females) aged between 18-40 years were recruited. Electrodermal activity (skin conductance level (SCL) and skin conductance response (SCR) amplitude) and Respiratory Sinus Arrhythmia (RSA) were assessed during the Emotional Stroop Task (EST). EST having expressive emojis superimposed with the congruent and incongruent words was used.

**Result:** Mean SCL and SCR amplitude was significantly increased during EST in incongruent and congruent blocks as compared to neutral block (respectively: 14.64 ± 6.73, 12.99 ± 6.26 vs. 7.75 ± 4.93 μS, p < 0.001 and 0.182 ± 0.168, 0.158 ± 0.134 vs. 0.021 ± 0.015 μS, p < 0.001). RSA was significantly decreased in incongruent and congruent blocks as compared to neutral blocks (respectively: 36.47 ± 10.53, 39.40 ± 10.15 vs. 48.66 ± 10.27 msec2, p < 0.001). We found an increased sympathetic activity and parasympathetic withdrawal while performing the task.

**Conclusion:** The results of this study suggested that emojis are adequate stimuli to elicit autonomic responses and change both sympathetic (EDA) as well as parasympathetic responses (RSA). Males and females showed similar autonomic arousal for emoji but the baseline emotional status was different for both genders.

## Introduction

The understanding of messages in digital communication can be affected by the lack of gestures and facial expression(1). To overcome this problem, the pictorial character having expression i.e. emojis, came into being. They not only ample the magnetism but also aid the emotional essence of the message(2). Emotions are produced by different neuronal networks in the brain associated with various neurophysiological changes in the body. These changes in the body and behavior of individuals are important for adaptation during various situations for survival. The brain perceives the experience for a particular emotion and conveys information to produce bodily sensations accordingly. The responses to emotions could vary among males and females due to their anatomical and hormonal disparity.

The amygdala plays a key role in emotional processing and emotional memory. The subnuclei of the amygdala receive emotional information from the cortical and various subcortical structures and give an output to the hypothalamus(3). Hypothalamus in turn activates the autonomic nervous system. Various neurophysiological studies suggest changes in electrodermal activity, heart rate, respiratory rate, and blood pressure during emotional arousal(4,5). These responses vary from individual to individual depending on the degree and intensity of the emotional stimuli. Autonomic responses are also influenced by the age and gender of the person. Previous studies indicated that there were distinct reactions in males and females to emotional primes(6). The difference in functional brain structures and limbic and paralimbic systems of males and females pronounced greater sensitivity in females to stressful emotional stimuli. A facial EMG study revealed that females are more facially reactive to happy and angry facial expression stimuli than males(7).

Unlike human facial expressions, emojis dominate in the virtual technical era to convey thoughts(8). Emojis have been used for conveying the contextual meaning of messages to receivers because their neural responses resemble face-to-face communication(9,10). So, it can be inferred that the use of emojis can also cause some autonomic changes in the human body. However, the changes in autonomic reactivity due to these emojis are still elusive.

The autonomic responses related to emotional stimuli such as face and words have been assessed by many researchers. Most of these studies were focused on electrodermal activity (EDA), which is used as a clinical measurement of emotional stress(11). EDA is the electrical activity of the eccrine sweat gland, innervated mainly by sympathetic postganglionic cholinergic fibers(12). Skin responses could be observed in 2 forms; tonic component and phasic component. The tonic component is determined by Skin conductance level (SCL). The short-term (phasic) changes in sympathetic activity are determined by skin conductance responses(SCR) which are produced by specific stimuli such as emotional stimulus (13,14). In addition, EDA is affected by the gender of the individual. Males have comparatively higher EDA than females due to the variation in morphology of sweat glands and the activity of sweat glands is higher(15).

Emotional stimuli also affect the heart rate and respiratory rate. The firing activity of the sinoatrial node is influenced by the fluctuating breathing pattern i.e. increases during inspiration and decreases during expiration(16), referred to as respiratory sinus arrhythmia (RSA)(17). RSA is used to measure the parasympathetic changes associated with emotions(18,19).. RSA is measured as the change in inter-beat interval during each respiratory cycle(20). It is phasic control of cardiac activity by the vagus nerve which increases during resting condition as parasympathetic activity dominates in resting condition (21–23).

EDA and RSA have been used widely to assess the alteration in sympathetic and parasympathetic activities related to emotions during neuropsychological tasks such as the Stroop task(24–26). The emotional Stroop task is the most commonly practiced tool in psychological studies, also used during the autonomic assessment to produce emotional stress(27).

The present study aimed to assess the changes in autonomic responses (EDA and RSA) to emotional stress induced by emojis in males and females. A modified emotional Stroop task (EST) having expressive emojis (happy, sad, fear, and angry) superimposed with precedent emotional words was designed to produce emotional arousal.

### Materials and methods

The study was conducted in the Cognitive Neurophysiology laboratory and initiated after approval by the institutional ethics committee, All India Institute of Medical Sciences Jodhpur. A total of 100 healthy volunteers (50 males and 50 females) were recruited after informed consent. The participants were in the age group of 18 to 40 years (mean ± SD - 27.87 ± 5.37 years). They were included based on education (up to 5^th^ standard), familiarity with the use of computers, and any history of disease affecting the autonomic nervous system (ANS) and respiratory system. The psychological general well–being index (PGWBI) was used to assess the emotional stability of individuals before their inclusion in the study. The participants having a raw index score (RIS) equal to or more than 73 were considered emotionally, physically, and mentally healthy and were included in the study.

### Procedure

Volunteers were asked to visit the lab between forenoon hours (09:00 am to 01:00 pm), to alleviate the effect of circadian influences, at the Cognitive Neurophysiology laboratory. The temperature of the lab was maintained in the thermo-neutral zone and the environment was noise-free to allay any undue anxiety. As soon as participants arrived in the laboratory, information about height, weight, gender, age, and hours spent on the internet was asked. The procedure of the study was verbally explained to the participants followed by a trial. Blood pressure was measured before and after the task. Participants were seated in front of the computer screens. EDA and ECG electrodes were attached and the respiratory belt was tied over the chest of the participant at the level of the fifth intercostal space. EDA and RSA parameters were recorded by using the Biopac MP150™ system (BiopacSystem, Inc. (C) Copyright 2000-2014, USA). For EDA recording, disposable silver/silver chloride disc electrodes (EL507, 3M, India Ltd.) along with the EL101 isotonic gel were used. BioNomadix™ Wireless EDA recording module was tied on the wrist and then connected with the electrode placed on their thenar and hypothenar eminences. A skin conductance module PPGED-R™ amplified electrical signals. Lead II ECG signals and respiratory movements have been recorded by RMS™ physiograph (Recorders and Medicare system (P) Ltd. / POLYG1311020) using a respiratory belt (RMS™). These systems were already connected to a signal pre-amplifier UIM100C™, which is a universal interface module, and this pre-amplifier in turn was connected to a computer equipped with Biopac MP150™ via Ethernet. Acqknowledge™ software version 4.4 was used to visualize the data in real-time and also store the data. The baseline ECG, respiratory rate, and skin conductance level were recorded for 5 minutes and were continued throughout the task. After the completion of 5 minutes, the Emotional Stroop Task containing emojis of different emotional valence was presented on the computer screen via Superlab5.0™ (Cedrus Corporation). The emotional Stroop task contains three blocks, a neutral block, a congruent block, and an incongruent block. Neutral block consists of the yellow circle only whereas in congruent block emojis have 4 prime emotions (happy, sad, fear, and angry) superimposed with similar emotional words (happy, sad, fear, and angry) as represented by particular emoji. Unlike congruent block, in incongruent block emotional words conflict with the emotions of emojis. Participant has to press the assigned key according to the facial emotion of emojis while ignoring the emotional words. To ensure attention, participants were asked to press the response key as soon as any emoji was presented on the screen. The task was synchronized with data collection.

EDA, ECG, and respiratory data were acquired at 2 kHz. RSA required two signals to perform the analysis – ECG Lead II and respiration. The difference between the maximum and minimum RR interval per breath measured the RSA values.

But to reduce the computational load, EDA data was resampled at 200 samples per second and analyzed(28). Phasic component (SCR amp) was obtained from SCL data using AcqKnowledge™4.4 software for the neutral, congruent, and incongruent blocks.

### Statistical Analysis

The data obtained was analyzed using SPSS ver.20 software (IBM). The data was found normally distributed on normality tests and therefore expressed as Mean ± SD. The statistical analysis was done using repeated measure ANOVA followed by post hoc Tukey’s test for comparison among all three blocks. Gender difference in autonomic responses was done using an independent sample t-test. P value ≤ 0.05 was considered as significant.

## Results

The study evaluated the autonomic responses to emotional Stroop task in 100 emotionally stable healthy volunteers in the age group of 18 – 40 years (27.87 ± 5.37 years) with an equal number of males and females. The autonomic responses (EDA and RSA) have been assessed for neutral, congruent, and incongruent blocks during EST and compared between males and females.

### 1. Electrodermal Activity (sympathetic nervous system activity)

The electrical conductance of skin was calculated for neutral, congruent, and incongruent blocks. The mean skin conductance level (SCL) was higher for the incongruent block (14.64 ± 6.73 μS) and congruent block (12.99 ± 6.26 μS) compared to the neutral block (7.75 ± 4.93 μS). SCL in all three blocks i.e. incongruent, congruent, and neutral in males (18.65 ± 6.20 μS, 16.52 ± 6.07 μS and 10.16 ± 4.99 μS respectively) was higher as compared to females (10.63 ± 4.50 μS, 9.46 ± 4.13 μS and 5.33 ± 3.52 μS respectively). Differences in SCL were statistically significant (p < 0.001).

**Table 1.1:** Showing the mean Skin Conductance Level for neutral, congruent, and incongruent blocks during the Emotional Stroop task (Mean ± SD)

| Skin Conductance Level (microSiemen) | Neutral block | Congruent block | Incongruent block | p-value |
| --- | --- | --- | --- | --- |
| Overall | $7.75 \pm 4.93$ | $12.99 \pm 6.26$ | $14.64 \pm 6.73$ | $< 0.001$ |
| Males | $10.16 \pm 4.99$ | $16.52 \pm 6.07$ | $18.65 \pm 6.20$ | $< 0.001$ |
| Females | $5.33 \pm 3.52$ | $9.46 \pm 4.13$ | $10.63 \pm 4.50$ | $< 0.001$ |

P value $\leq 0.05$ was considered as significant.

**Figure 1.1.**
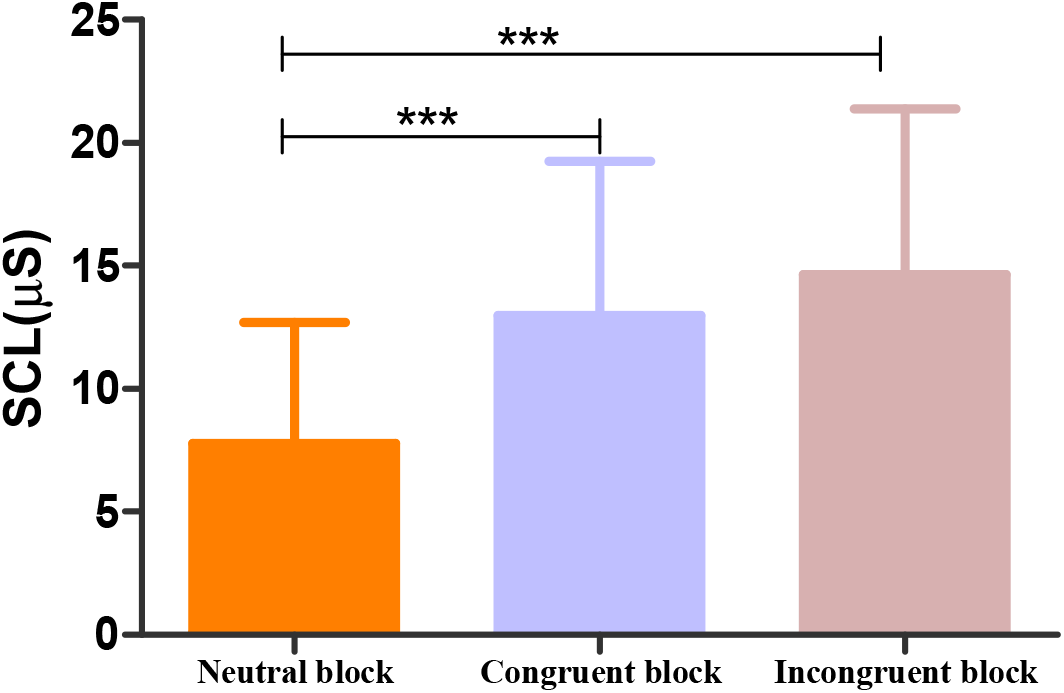
Column bar graph showing the difference between skin conductance levels (SCL, microSiemen (μS) in neutral block (NB), congruent block (CB), and incongruent block (InCB) during Emotional Stroop Task. The horizontal line of each plot represents the mean values of each parameter; the upper hinges correspond to the standard deviation (SD). A significant difference between the congruent block^***^ and the neutral block, and also between the incongruent block^***^ and the neutral block (*** p < 0.001).

The increase in skin conductance level of more than 0.02 μS was considered as a specific response. These specific responses were measured as the skin conductance response amplitude (SCR amp.). Average SCR amp. was higher for the incongruent block (0.182 ± 0.168 μS) and congruent block (0.158 ± 0.134 μS) compared to the neutral block (0.021 ± 0.015 μS). SCR amp. for males (0.238 ± 0.204 μS, 0.193 ± 0.156 μS, and 0.023 ± 0.014 μS) was higher as compared to the females (0.126 ± 0.096 μS, 0.122 ± 0.096 μS and 0.019 ± 0.017 μS) for all three blocks i.e. incongruent block, congruent block, and neutral block respectively. The data was statistically different between 3 blocks (p < 0.001).

**Table 1.2:**
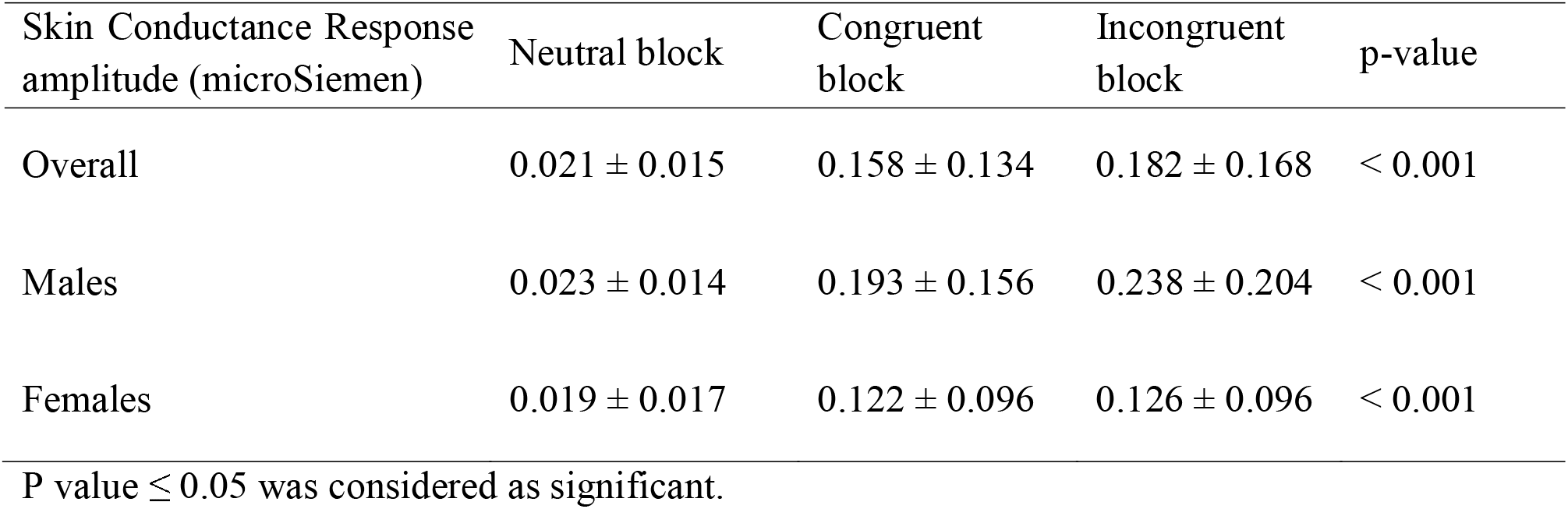
Showing the mean Skin Conductance Response amplitude for neutral, congruent, and incongruent blocks during the Emotional Stroop task (Mean ± SD).

**Figure 1.2.**
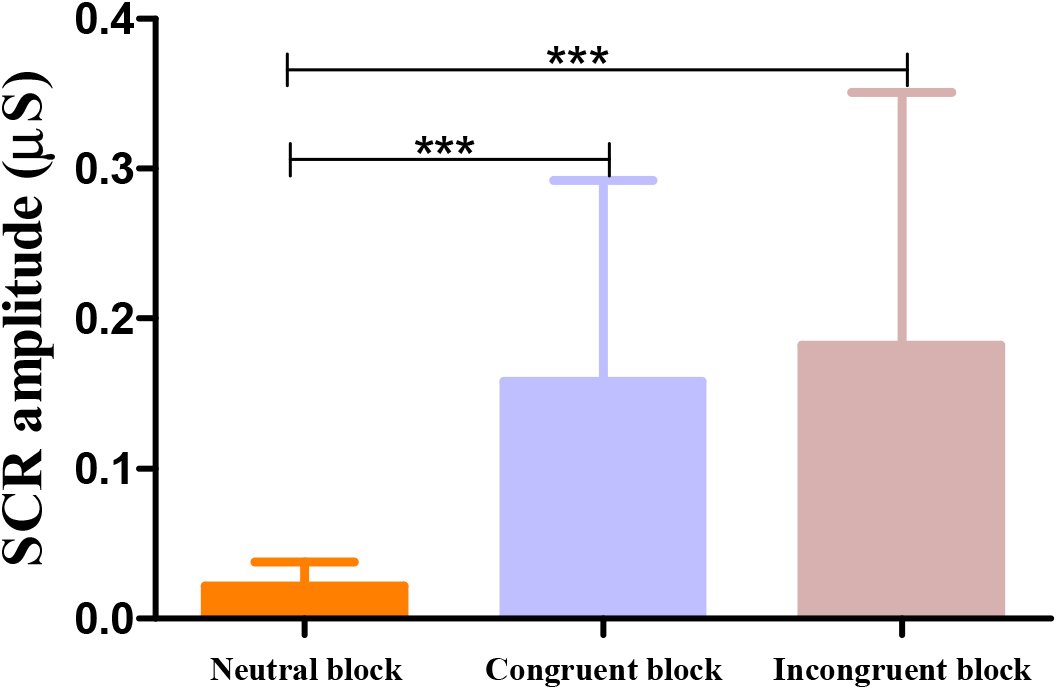
Column bar graph showing the difference between skin conductance response amplitude (SCR amp., microSiemen (μS) in neutral block (NB), congruent block (CB), and incongruent block (InCB) during Emotional Stroop Task. The horizontal line of each plot represents the mean values of each parameter; the upper hinges correspond to the standard deviation (SD). A significant difference between the congruent block^***^ and the neutral block and also between the incongruent block^***^ and the neutral block (*** p < 0.001).

Additionally, the absolute change and percentage change were calculated and compared between the two genders. The result of the independent sample t-test indicated that the difference between the two genders in absolute change in electrodermal activity parameters during EST was statistically significant (the SCL congruent condition [t (98) = 3.755, p < 0.05], incongruent condition [t (98) = 5.139, p < 0.05] and for SCR amp condition [t (98) = 2.619, p < 0.05], incongruent condition [t (98) = 3.368, p < 0.05]). Whereas, males and females were similar in the percentage changes for electrodermal activity parameters (SCL and SCR amp.) in congruent and incongruent blocks. This suggests that though the sympathetic responses to emoji are different in the two genders, the amplitude of the response is similar in them. This again reiterates that emojis have the power to evoke sympathetic responses similar in males and females.

**Figure 1.3.**
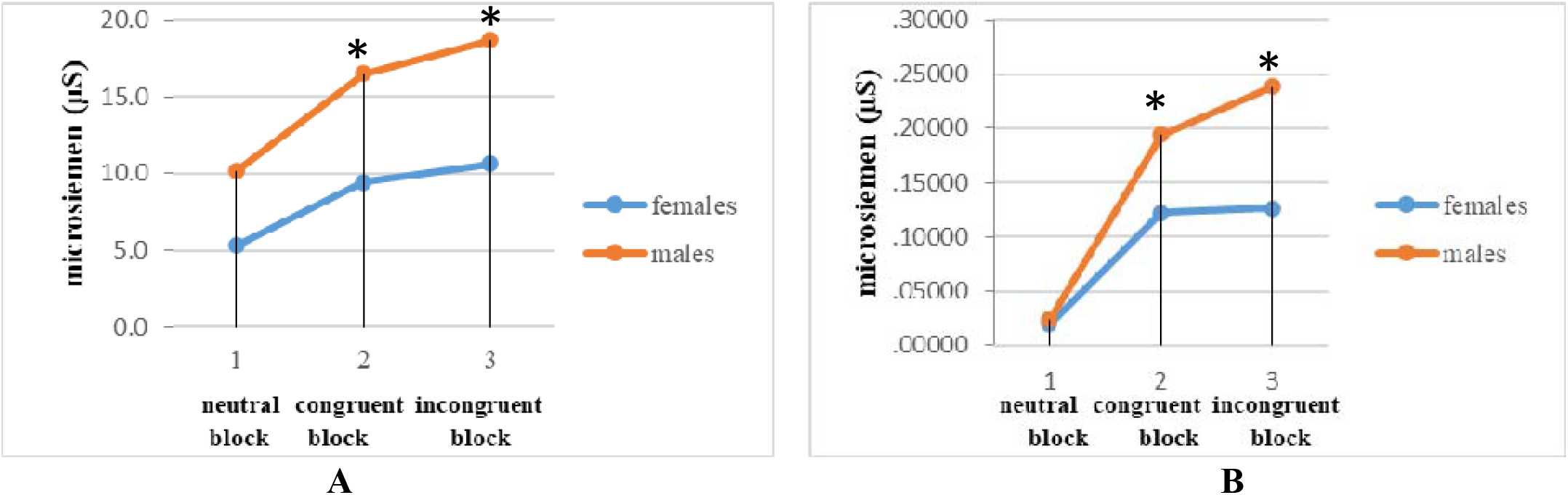
The average value of electrodermal activity parameters among males and females while performing the EST. The vertical dashed line indicates the beginning of a neutral, congruent, and incongruent block of EST. (A) skin conductance level (μS). (B) skin conductance response amplitude (μS). (* P < 0.05)

### 2. Respiratory Sinus Arrhythmia (RSA) – parasympathetic nervous system activity

For the calculation of RSA, the difference in maximum to minimum heart period was calculated for neutral block, congruent block, and incongruent block. The mean RSA was found to be decreased during incongruent (36.47 ± 10.53 msec^2^) and congruent block (39.40 ± 10.15 msec^2^) as compared to neutral block (48.66 ± 10.27 msec^2^). The average value of RSA for males was lesser (35.72 ± 12.17 msec^2^, 37.95 ± 12.18 msec^2^ and 49.72 ± 12.90 msec^2^) as compared to the females (37.74 ± 9.88 msec^2^, 41.37 ± 10.17 msec^2^ and 50.82 ± 10.12 msec^2^) for all three blocks i.e. incongruent, congruent and neutral respectively. The difference in RSA was statistically significant (p < 0.001).

**Table 2:** Showing the mean Respiratory Sinus Arrhythmia for neutral, congruent, and incongruent blocks during the Emotional Stroop task (Mean ± SD).

| Respiratory Sinus Arrhythmia<br>(millisecond square) | Neutral block | Congruent<br>block | Incongruent<br>block | p-value |
| --- | --- | --- | --- | --- |
| Overall | 50.27 ± 11.55 | 39.66 ± 11.29 | 36.73 ± 11.08 | < 0.001 |
| Males | 49.72 ± 12.90 | 37.95 ± 12.18 | 35.72 ± 12.17 | < 0.001 |
| Females | 50.82 ± 10.12 | 41.37 ± 10.17 | 37.74 ± 9.88 | < 0.001 |

P value ≤ 0.05 was considered as significant.

**Figure 2.1.**
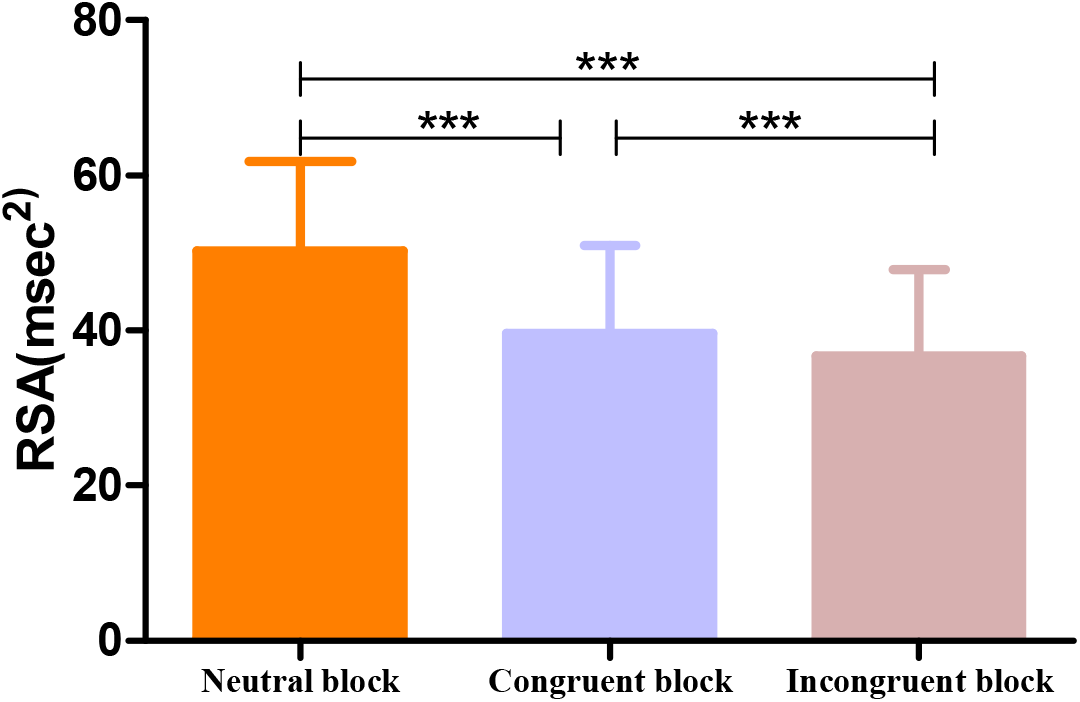
Column bar graph showing the difference between respiratory sinus arrhythmia (RSA, milli second^2^) in neutral block (NB), congruent block (CB), and incongruent block (InCB) during Emotional Stroop Task. The horizontal line of each plot represents the mean values; Upper hinges correspond to SD. A significant difference between all three blocks (*** p < 0.001).

On further analysis for gender, the result of the independent sample t-test showed the difference in absolute change and the percentage change of RSA value among males and females was statistically not significant (p > 0.05). Though the combined data of males and females showed that emojis can elicit parasympathetic responses when compared between the genders, we could not find any difference. This could be explained by a small amount of change observed.

**Figure 2.2.**
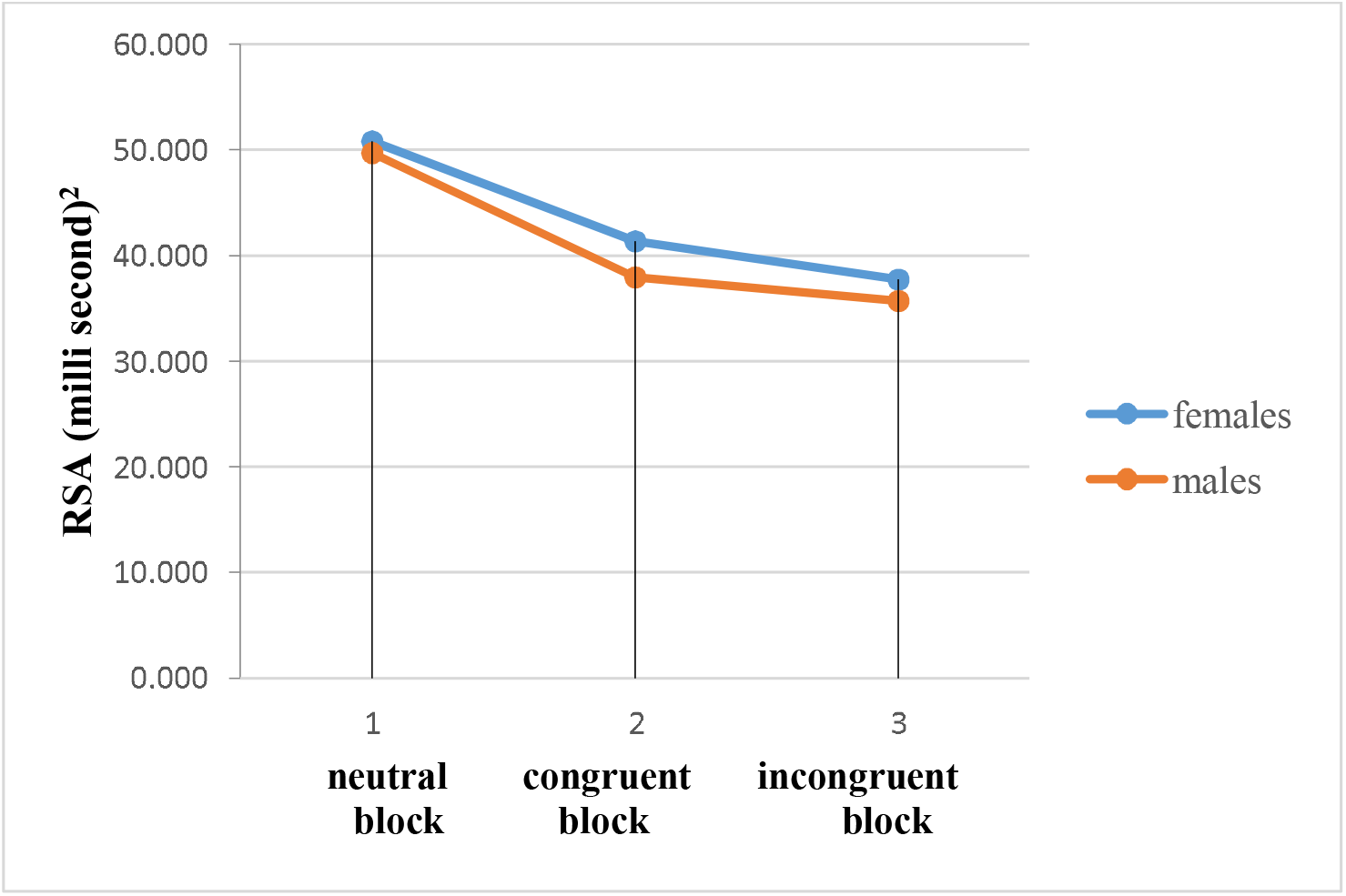
The average value of respiratory sinus arrhythmia (msec^2^) among males and females while performing the EST. The vertical dashed line indicates the beginning of the neutral, congruent, and incongruent block of EST.

## Discussion

The present study assessed the autonomic parameters (EDA and RSA) of 100 healthy subjects (50 males and 50 females) during emoji-word EST among neutral, congruent, and incongruent block images. The result showed that sympathetic activity was increased during EST. Sympathetic activity was found higher for incongruent as well as congruent blocks which contained emojis compared to the neutral block. Parasympathetic activity showed an opposite trend as RSA was found to be minimum for incongruent block and maximum for neutral block showing the greater parasympathetic dominance during neutral block which does not produce any emotional stress. The results were consistent with previous studies.

There was a significant increase in skin conductance level (SCL) and skin conductance response (SCR) amplitude for both incongruent block and congruent block having both emotional words and expressive emojis (p < 0.001). The greater sympathetic activity for the incongruent block compared to the congruent block was probably due to the increased stress level and attention of volunteers during the incongruent block compared to the congruent block. The results also showed minimum sympathetic activation during the neutral block which indicates that emojis served as adequate stimuli to alter the autonomic nervous system activity. The strength of evoked emotion corresponds to the magnitude of SCR, so magnitude can be used as a quantitative marker of emotional arousal(29). Emojis are used in digital communication whereas emoticons are used for textual language. Thompson et al. in their study used the written language with or without emoticons to assess the responses using electrodermal activity and facial electromyography. They reported an increased electrodermal activity for messages having emoticons along with an increase in mean EMG amplitude (i.e. mean EMG corrugator activity) due to the increased smiling. The authors conclude that emoticons elicit autonomic responses and modulate the emotional impact of messages(30) is similar to our findings in the present study. In contrast, Ruiz-Robledillo et al. found lower electrodermal activity to acute stress in caregivers of people with ASD than non-caregivers. They suggested the probable reason for this hyporeactivity with the adaptive habituation to stress due to long-term exposure in the caregivers that could protect their health(31).

On in-depth analysis using gender as a grouping variable, there was a significant increase (p < 0.001) in skin conductance level (SCL) and skin conductance response (SCR) amplitude during emotional Stroop task in males as compared to females. Martinez-Selva et al. also evaluated thes gender and menstrual cycle difference in electrodermal activity. They reported that the males had greater electrodermal activity as compared to females. In female, the preovulatory females had augmented values as compared to postovulatory. The possible reason of this difference could be the greater sweat gland activity in males compared to females (6,10)(33).

We obtained a significant decrease in RSA progressively from neutral block to congruent to incongruent block during the EST which showed greater withdrawal of parasympathetic activity during incongruent block images (p < 0.001). The reason for this decrease in RSA from its baseline could be the emotional arousal produced by emojis that allowed heart to beat faster as the parasympathetic control was withdrawn. The change in RSA could be appreciated, when healthy individuals are acquainted with the movie clip or kind of task like Stroop task having emotional content. Exposure to emotional stress can cause parasympathetic withdrawal which allowed heart to beat faster and at the same time respiratory rate also increases, so RSA decreases. ASD patients showed lesser decrease in RSA during EST than the healthy individuals which probably could be due to the habituation to stress (34).

RSA among males and females showed a no significant difference in our findings. The average RSA was found to be higher in females compared to males during emotional induction caused by emojis. The parasympathetic withdrawal was found more in males than females that showed the males have greater RSA than females(35). The probable reason of this difference may be due to difference in anatomy of respiratory structures and also difference in lung volumes (such as tidal volume). Males have more tidal volume than females. Additionally, RSA is inversely proportional to respiratory rate i.e. RSA decreases with increase in respiratory rate and vice-versa(36). Another proposed reason for this difference is inhibitory effect of prefrontal cortex on sympathoexcitatory subcortical circuits. This inhibitory effect was more in females as indexed by HRV and heart rate(37). In contrast to our study, Feurer et al. obtained a greater decreased RSA in girls compared to boys living in a neighborhood having exposure of crime in middle childhood. The results indicated that girls living in neighborhood which had high risk of violent crimes (emotional stimulus) showed decreased RSA compared to boys living in the same neighborhood. The authors attributed this difference to the stronger activation of cortical and limbic structures in females for angry and fearful stimuli in comparison to males(38). Another reason for this difference could be the age of the children. As reported earlier children have higher overall HRV as compared to Adults and girls have lower values of HRV than boys.

There are some limitations to our study, small sample size and assessment of autonomic responses for each valence of emotion. It was not feasible to complete the data collection more than 100 volunteers (without error) on short study period of one year. Also, taking into account the valence of emotion needs an elaborated methodology and refined assessment tools which were difficult to incorporate in study design due to time and resource constraints.

## Conclusion

We can conclude that emojis do modulate the autonomic activity when presented in the form of emotional stimuli by increasing the sympathetic arousal and withdrawing the parasympathetic activity. Our finding suggests that though male female are different in their baseline emotional status but their autonomic responses to the emojis were similar. Since characteristic autonomic responses are features of emotional state, we can surmise that emojis are representative of emotion, heighten arousal, clarify intentions and potentially aid comprehension equally in both genders.

As the use of emojis can alter the autonomic responses, it imply that overuse of emojis can be deleterious to mental as well as physical health. These findings may provide a basis to restrict the use of social media in patients those suffering from conditions like myocardial infarction (MI). Alteration of autonomic activity due to overuse of emojis may be detrimental in patients with autonomic disorders.

## Acknowledgment

We wish to thank Dr. Shival Srivastav, Assistant Professor, Department of physiology, AIIMS, Jodhpur who supported us during the conduction of the study. We would like to show our gratitude to all the participants without whom the study could not have been possible.

