## Supplementary figures and images for "Effect of Emoji on Autonomic Nervous System: Evidence from EDA and RSA studies"

### Supplement Figure 1

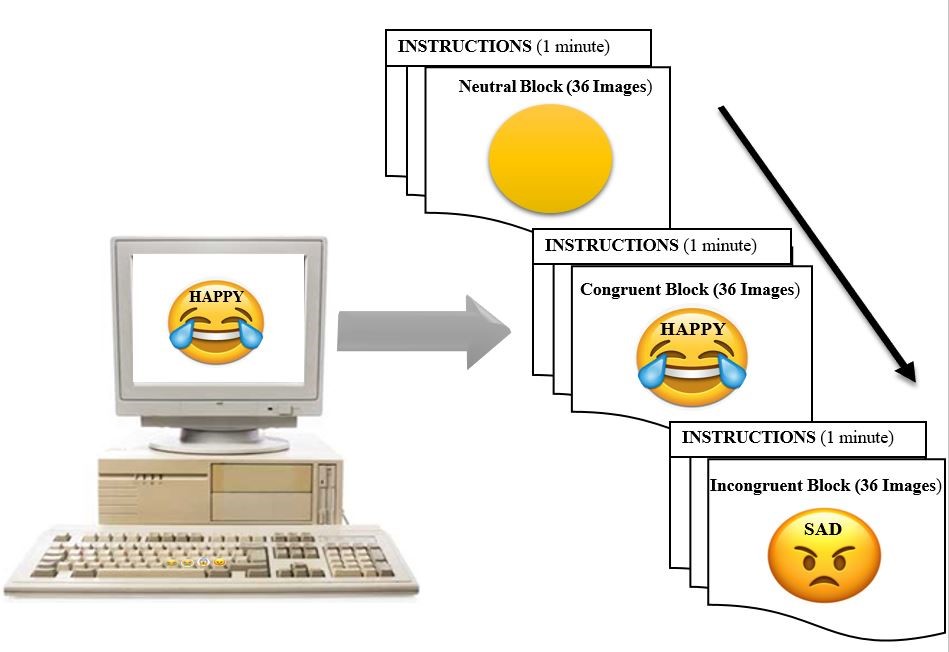
